# Spatial frequency channels mediate a mental ruler for spatial vision in humans

**DOI:** 10.1101/2024.10.15.615126

**Authors:** Shijia Zhang, Ce Mo, Lei Mo, Fang Fang

**Author notes:** equal contribution (Shijia Zhang, Ce Mo).

## Abstract

An intriguing mystery concerning spatial vision is that the perceived spatial property information of the same stimulus varies in different contexts. While previous findings are suggestive of an adaptable mental ruler by which the raw retinal signals are flexibly scaled according to the prevailing context, whether and how such a putative mental ruler is utilized by the visual brain remain largely unknown. We hypothesized that the putative mental ruler was represented by multiple differently-tuned spatial frequency (SF) channels. A finer division of the ruler, signaled by dominance of the high-SF channels, is translated to a wider-spread topography of neural activation via concentration of neuronal receptive fields, causing perceptual inflation. Combining psychophysics and fMRI, we found that modulation of SF channels caused a systematic distortion in perceived separation, a representative fundamental spatial property, and a global displacement of population receptive fields (pRF) in primary visual cortex. Computational modeling further showed that both the perceptual distortion and the global pRF displacement were functionally coupled with the magnitude of SF channel modulation. Our findings reveal, for the first time, an adjustable mental ruler that commonly governs the perception of spatial property information in different contexts, and suggest a cognitive scaling system based on SF channel reweighting and displacement of neuronal RFs.

## Introduction

The core of spatial vision concerns the extraction of the spatial property information (e.g., size, separation, etc.) of visual objects from the light signal patterns impinging on the retinae (De Valois & De Valois, 1988). However, our spatial vision is rarely a veridical Cartesian projection of the external physical world, but rather is shaped by different contexts. As such, the raw retinal signals must be processed in a specific manner to generate the perceived spatial property information that are appropriate to the prevailing visual context (Sperandio & Chouinard, 2015). An intriguing idea derived from previous studies (Dakin et al., 2011; Hisakata et al., 2016; Parkinson et al., 2014; Walsh, 2003) is that this process is governed by an adaptable mental ‘ruler’ whose division could be flexible adjusted in response to contextual modulation. A finer division of the mental ruler causes the same visual object to span more ruler graduations, thereby leading to an inflation in spatial perception, while a coarser division causes a shrinkage. At the neural level, however, distortions in spatial property information perception have been attributed to the topographic profile of neural activation across early visual cortex (Ho & Schwarzkopf, 2022; Moutsiana et al., 2016; Sperandio et al., 2012). For example, the spatial spread of neural activation in primary visual cortex (V1) elicited by a visual stimulus closely matches the perceptual, rather than the retinal, stimulus size altered by adaptation (Pooresmaeili et al., 2013) or by depth cues (Fang et al., 2008; Murray et al., 2006). As a result, whether such a putative mental ruler is represented and utilized by the visual system remains an open question, because it is not understood how the putative mental ruler is translated to the topography of neuronal activities.

The translation from the putative mental ruler to the neural activation topography is likely contingent on a bank of sensory pathways that are tuned to different spatial frequencies (SF), i.e. SF channels. On one hand, SF tuning is intrinsically related to the tiling of neuronal receptive fields (RF) that plays a decisive role in shaping the topographic profile of neural activities (Altan et al., 2024; He et al., 2015; Ni et al., 2014). Dense tiling of neuronal RFs is accompanied by the population’s tuning preference of high SFs, as it allows for the resolution of small differences of local contrast patterns elicited by spatially adjacent stimuli (Aghajari et al., 2020; Anton-Erxleben & Carrasco, 2013). On the other hand, sensitivity to different spatial scales is constrained by the relative weighting of the interacting SF channels (Sowden & Schyns, 2006). Enhancing the high-SF channels sharpens the population’s representation of fine-grained stimulus information, while biasing towards the low-SF channels instead optimizes the processing of coarse-grained information (Flevaris & Robertson, 2016). We therefore hypothesized that the putative mental ruler was represented by multiple differently-tuned SF channels, with the smallest division corresponding to the inverse of the peak of the population SF response (i.e. the area of one luminance variation cycle). Specifically, the dominance of high SF channels signals a finer division of the ruler, which is translated to a wider-spread neural activation topography via concentration of neuronal receptive fields, thereby causing an inflation in spatial perception (**Figure 1**).

**Figure 1.**
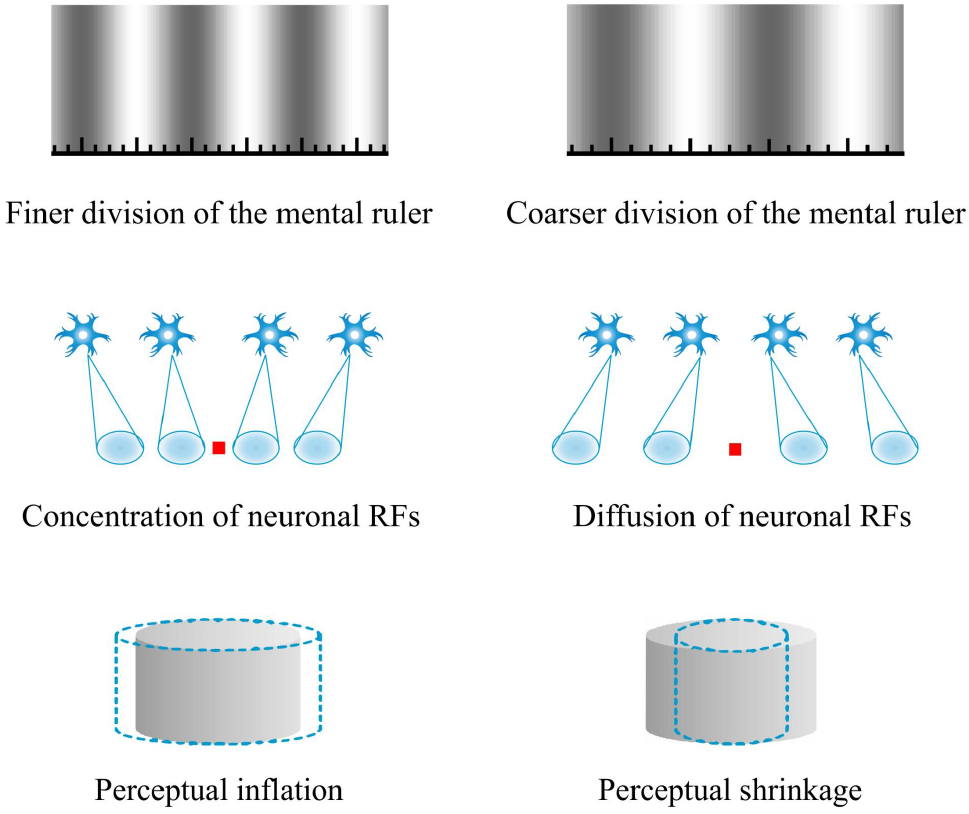
Demonstration of the mental ruler hypothesis. The putative mental ruler (shown as bold black line segments inset) with a finer and a coarser division is signaled by the dominance of the high-SF channels **(Top left)** and the low-SF channels **(Top right)**, which is translated to the neural activation topography via a concentration **(Middle left)** and a dispersion **(Middle right)** of neuronal RFs (shaded ellipses), respectively. A finer division of the mental ruler causes the same physical stimulus to span more ruler graduations, which inflates the perceptual readout (dashed contour) **(Bottom left)**, while a coarser division causes a perceptual shrinkage **(Bottom right)**.

The key to examining our hypothesis is to investigate whether modulation of these SF channels displaces the neuronal RFs in early visual cortex and alters spatial property perception. Here, we employed the perception of separation, a representative fundamental spatial property, as our test bed. Using psychophysical adaptation, we found that the perceived separation, whether measured directly or indirectly, was consistently reduced by inhibition of the high-SF channels in comparison to inhibition of the low-SF channels. Conversely, driving these channels produced an opposite effect. At the neural level, the perceptual distortions induced by SF channel modulation aligned with a global displacement of the V1 population receptive fields (pRF). Finally, a computational model showed that both the perceptual distortion and the pRF displacements were coupled with the magnitude of SF channel modulation. Together, these findings provided converging evidence for an adjustable mental ruler in spatial vision, and suggested a unified cognitive scaling mechanism based on SF channel reweighting and displacement of neuronal RFs.

## Results

### SF channels modulation via adaptation distorts perceived separation

We first investigated whether perception of separation was influenced by selective modulation of SF channels using the psychophysical adaptation technique. Adaptation is a time-honored tool in psychophysics, which leverages a general property of sensory neurons that prolonged exposure to a stimulus feature selectively reduces the contribution of the underlying neuronal population tuned to that feature. In the psychophysical adaptation experiment, subjects simultaneously adapted to two sinusoidal gratings with different SFs that were drifting in synchrony at the same speed (**Figure 2a**). One of the gratings had a constant SF of 0.25 cycles/° (fixed adaptor), while the SF of the other grating varied from 0.25 to 1.5 cycles/° (matching adaptor) in different blocks. This design was intended to maximize the sensitivity to the SF channel modulation effects. After adaptation, subjects performed two different tasks on the stimuli that were presented at the same locations as the adaptors. In the dot-pair task, the task stimuli were two pairs of dots and subjects reported the pair with a larger apparent gap. In the shape task, the stimuli were two ellipses with their major axes aligned vertically and subjects reported the ellipse that appeared more elongated (**Figure 2b**). Note that the shape task essentially engaged in comparing the perceived separation between the endpoints of the two ellipses’ minor axes. In both tasks, the task stimulus ipsilateral to the fixed adaptor served as the standard stimulus that remained invariant throughout the experiment, while the task stimulus ipsilateral to the matching adaptor (or equivalently, contralateral to the fixed adaptor) served as the probe that was varied from trial to trial using a staircase procedure.

**Figure 2.**
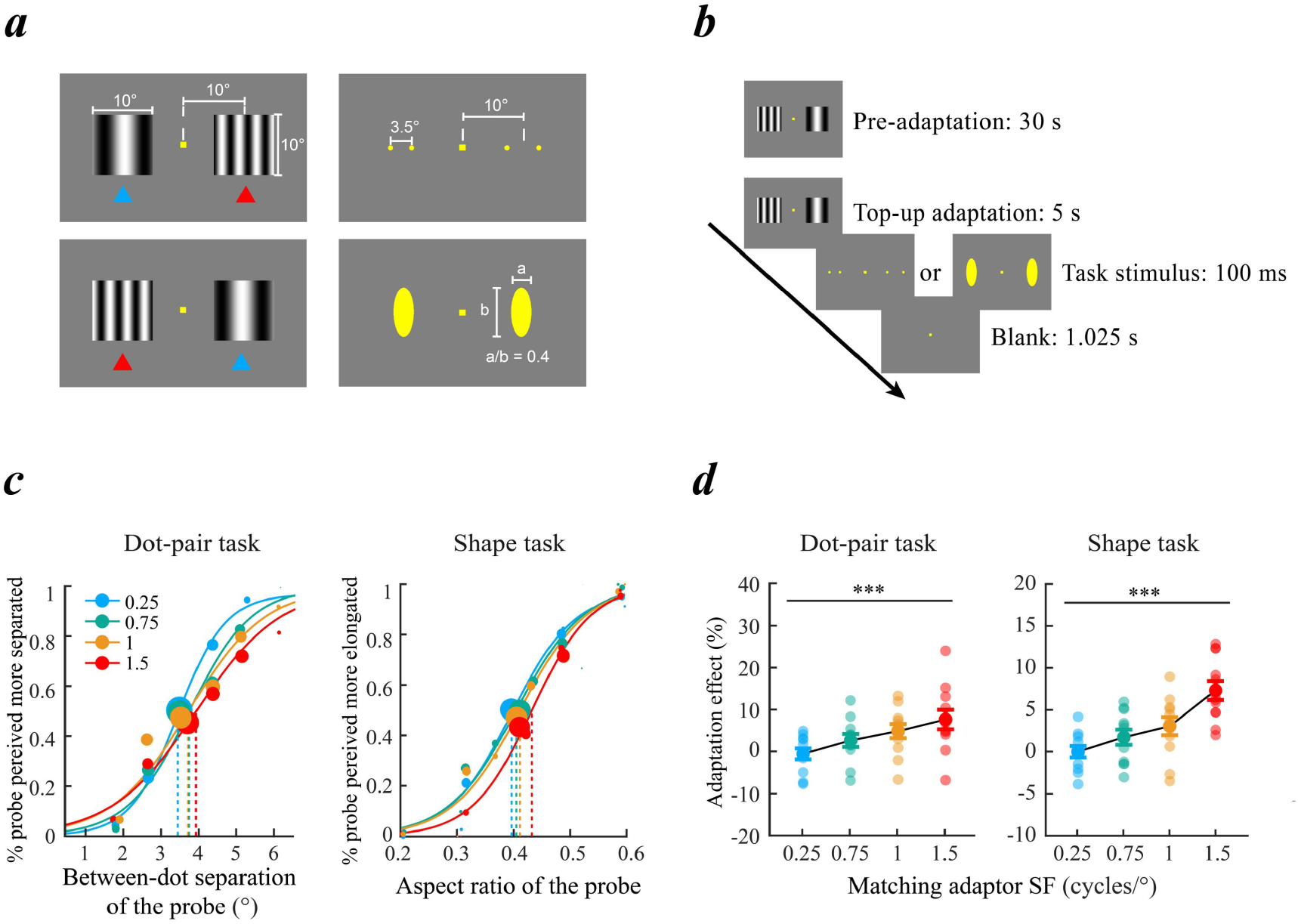
Psychophysical adaptation experiment. Experimental stimuli. **Left)** Illustration of the adapting stimuli. Both the fixed adaptor (indicated by the blue arrowhead) and the matching adaptor (red arrowhead) could appear in either visual field. **Right)** Illustration of the task stimuli. In this example, the standard stimulus of the dot-pair task (upper) is shown in the left visual field, while that of the shape task is shown in the right visual field. The standard stimuli were always preceded by the fixed adaptor. ***b***. Experimental procedure of a staircase block. Subjects indicated the perceptually more separated pair of dots (dot-pair task) or the perceptually more elongated ellipse (shape task). ***c***. Psychometric curves corresponding to the four matching adaptors (coded in different colors) in each task, which were based on the data pooled from all subjects for visualization, with the size of the dot point proportional to the number of trials in that bin. Note that statistical analyses were conducted on individual fits. The abscissa refers to the gap (dot-pair task) or the aspect ratio (shape task) of the probe. The ordinate refers to the percentage of trials in which the probe was perceived more separated or more elongated. Dotted lines indicate the physical separation or aspect ratio corresponding to the 50% probability that subjects chose the probe over the standard stimulus (i.e. point of subjective equality of the standard stimulus, PSE). ***d***. Magnitude of the SF adaptation effect as a function of the matching adaptor SF in both tasks. Each point denotes the data from one subject in one matching adaptor condition. Error bars denote one standard error of the mean (S.E.M.) across all subjects. **: *p* < 0.01. ***: *p* < 0.001.

In both tasks, the behavioral data showed a similar pattern regardless of the visual field in which the fixed adaptor was presented and were thus pooled together for analysis. For each matching adaptor, the data were fitted with a Logistic function and we obtained the psychometric curve that described the percentage of trials in which the probe was chosen as a function of the between-dot gap (dot-pair task) or the aspect ratio (shape task) of the probe. For the dot-pair task, when both adaptors shared the same SF (0.25 cycles/°), the psychometric curve centered at the gap of the standard stimulus. However, as the SF of the matching adaptor increased, the psychometric function showed a consistent horizontal rightward shift (**Figure 2c** left), indicating that the perceived gap became smaller as subjects adapted to increasingly higher SFs. A similar pattern was also found for the shape task (**Figure 2c** right). It appeared as if both types of probes were ‘squeezed’ and ‘stretched’ after adaption to the high and the low SFs, respectively. To quantify the SF adaptation effect for each matching adaptor, we identified the critical point at which the probe and the standard stimulus were perceptually indistinguishable (the point of subjective equality, PSE), and computed the relative difference between the PSE and the standard stimulus in percentage units. As shown in **Figure 2d**, when both adaptors shared the same SF, the adaptation effect did not significantly differ from 0 in either task (dot-pair task: mean ± S.E.M: -0.77 ± 1.23%, one-sample *t*-test: *t*_11_ = -0.62, *p* = 0.546; shape task: mean ± S.E.M: 0.13 ± 0.65%, one-sample *t*-test: *t*_11_ = 0.20, *p* = 0.845). In contrast, the adaptation effect exhibited a monotonically climbing pattern with the increasing matching adaptor SF, which was consistently observed in both tasks (dot-pair task: *F*_(3,33)_ = 9.37, *p* < 0.001, *η*^*2*^ = 0.460; shape task: *F*_(3,33)_ = 16.08, *p* < 10^-4^, *η*^*2*^ = 0.594). Notably, this effect could not be attributed to the difference in internal noise, as there was no significant difference in the slope of the psychometric curves across the matching adaptors in either task (one-way repeated-measure ANOVA: dot-pair task: *F*_(3,33)_ = 0.78, *p* = 0.515; shape task: *F*_(3,33)_ = 0.75, *p* = 0.533). These findings showed that modulation of different SF channels via adaptation led to a clear discrepancy in perceived separation regardless of whether it was measured directly or indirectly, which agreed well with our hypothesis.

### SF channel modulation is coupled with a global-scale displacement of pRFs in V1

We then investigated whether the spatial profile of neuronal RFs was influenced by modulation of SF channels. A well-established non-invasive approach of investigating neuronal RF profiles in human visual system is the fMRI-based population receptive field (pRF) mapping (Dumoulin & Wandell, 2008; Wandell & Winawer, 2015). Although the nature of fMRI data denies direct investigation of single neurons, the pRF mapping technique provides an effective, albeit circumstantial, characterization of neuronal RF profiles by modeling the joint receptive field of the underlying neuronal population within a voxel. However, since the neurons of interest are driven by the stimulus in pRF mapping, we needed to determine whether the perceptual effects induced by SF channel enhancement were opposite to those induced by SF channel inhibition, which would test our hypotheses from a complementary angle. Moreover, our findings from the psychophysical adaptation experiment were also constrained by two concerns. First, because the two adaptors were drifting at the same speed, different SF channels might experience stronger adaptation effects as they have different motion sensitivity profiles. Second, because the two adaptors were vertically orientated gratings, they might bias specific orientation channels over the others, making them less efficient for interrogating the SF channels. To address these issues, an out-scanner experiment was conducted in prior in which we explicitly validated the hypothesized perceptual distortion effect induced by SF channel enhancement without the involvement of orientation-specific information or stimulus motion.

In the out-scanner experiment, subjects were simultaneously presented with two oblique yellow bars superimposed on two square-shaped checkerboard backgrounds with different SFs. Based on the observations from the psychophysical adaptation experiment, we focused on the comparison between 0.25 cycles/° (low-SF background) and 1.5 cycles/° (high-SF background). It should be noted that the chromatic bars were employed to increase the saliency of task stimuli on the checkerboard backgrounds, such that the potential demands for covert attention, if any, were minimized. Similar as in the psychophysical adaptation experiment, the bar on the low-SF checkerboard served as the standard stimulus, while the bar on the high-SF checkerboard served as the probe that was varied by a staircase procedure (**Figure 3a**). Subjects indicated the bar that appeared longer, which was essentially a variant version of the dot-pair task. A psychometric curve was fitted to the behavioral data and we identified the PSE of the standard stimulus from the fitted curve. We found that the psychometric function was horizontally shifted to the left of 3.5°, i.e. the physical length of the standard stimulus (**Figure 3b** left), indicating a clear perceptual distortion that the bar on the high-SF background appeared longer than the bar on the low-SF background. The magnitude of the perceptual distortion effect was further measured as the relative difference between the standard stimulus and the PSE in percentage units, which was significantly above zero (mean ± S.E.M.: 3.8 ± 0.79%, one-sample *t*-test: *t*_*17*_ = 6.14, *p* < 10^-4^, Cohen’s *d* = 1.447) (**Figure 3b** right). These results showed that the enhancement of different SF channels also distorted perceived separation, producing a perceptual effect opposite to that induced by SF adaptation. Moreover, the perceptual distortion could not be explained by orientation contingency or difference in adaptation between SF channels, because there was no orientation-specific information or motion component in the checkerboard backgrounds.

**Figure 3.**
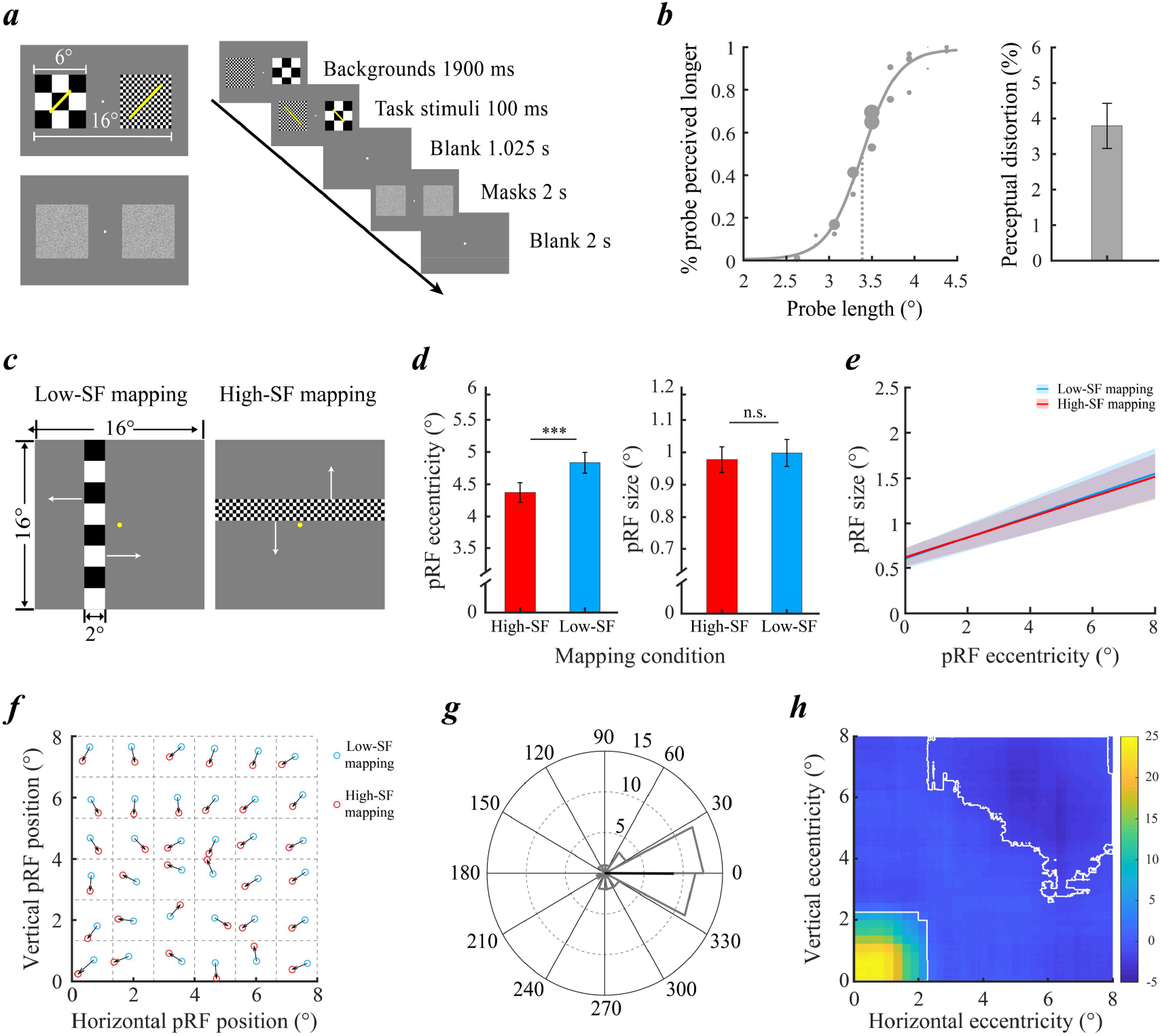
fMRI experiment. a. Design of the out-scanner experiment. **Left)** Illustration of the task stimuli (upper) and the Gaussian white noise masks (lower). The task stimuli were two oblique bars (the standard stimulus and the probe) superimposed on two different SF backgrounds (0.25 and 1.5 cycles/°). The standard stimulus was always superimposed on the low-SF background (presented in the right visual field in this example). **Right)** Single trial procedure. Subjects indicated which of the two bars appeared perceptually longer. ***b***. Psychophysical results. **Left)** the psychometric function fit to the binned data from all subjects for visualization. Size of the date point was scaled in proportion to the number of trials in that bin. Statistical analyses were conducted on individual fits. The dashed line indicates the PSE of the standard stimulus. **Right)** Magnitude of the perceptual distortion effect. Error bar denotes one S.E.M. across subjects. ***c***. Illustration of the high-(left) and the low-SF (right) mapping stimuli used for pRF mapping. White arrows indicate the possible sweeping directions of the bar stimuli. ***d***. Comparison of the pRF eccentricity and the pRF size of the included V1 voxels between the two SF mapping conditions. Error bars denote one S.E.M. across all subjects. ***: *p* < 0.001, n.s.: not significant. ***e***. Relationship between pRF size and pRF eccentricity in the two SF mapping conditions. The solid lines denote the linear fits across subjects. The shaded bands around the lines indicate the 95% confidence interval estimated using a bootstrapping method. ***f***. Average displacement vectors (across two binning schemes) pertaining to different parts of the visual space. Grey dashed lines denoted the boundaries of the grid cells that partitioned the merged quadrant. The blue circles denote the average of the mean pRF coordinates in the low-SF mapping condition, and the red circles denote the average of the mean pRF coordinates in the high-SF mapping condition. The norms of these vectors are scaled for better visualization. ***g***. Distribution of the angular deviations between the average displacement vector and the direction of the fovea in all grid cells. Specifically, a value of 0° indicates that the displacement directed at the fovea. The thick solid bar indicates the mean of the angular deviations. ***h***. Spatial map of the pRF density difference between the two conditions. White curves indicate the boundaries of the significant clusters identified using the non-parametric cluster-based analysis (*p* < 0.05, multiple comparisons corrected).

Having validated the perceptual distortion elicited by SF channel enhancement in the out-scanner experiment, we investigated whether and how it might alter the spatial profile of pRFs in early visual areas based on the same group of subjects. Each voxel-wise pRF was modelled as an isotropic 2-D Gaussian function. By stimulating different parts of the visual field with a mapping stimulus, we estimated the location and the size parameters of the Gaussian that yielded the best fit to the empirical BOLD response evoked by the mapping stimulus. The pRF mapping stimuli were the same checkerboard stimuli as in the out-scanner experiment, which were presented in an oriented rectangular aperture. The rectangular aperture swept through the entire visual field in multiple steps with a step duration of 2s (**Figure 3c**). Note that the step duration was the same as the total duration of checkerboard background presentation in the out-scanner experiment, hence the SF channels were driven by the same stimulus for the same duration in each step (i.e. 2s, see **Methods** for details) of the sweep. Each voxel-wise pRF was mapped using two otherwise identical stimulus sequences that differed only in the SF of the mapping stimulus (low-SF mapping condition: 0.25 cycles/°; high-SF mapping condition: 1.5 cycles/°). We focused on V1 as our primary region of interest due to its well-established role in processing spatial property information, as well as its highest sensitivity to changes in RF profiles afforded by the smallest RF size (Ho & Schwarzkopf, 2022; Schwarzkopf et al., 2011; Song et al., 2015). To avoid statistical bias in voxel selection, all the reported findings below were based on the same set of voxels in which the pRF model could explain at least 25% of the variance of the raw data in both SF mapping conditions (mean percentage of voxels included ± S.E.M.: 40.43 ± 2.12%). Notably, similar findings were obtained when we repeated the same analyses with two different criteria for inclusion of voxels (percentage of raw data variance explained ≥ 20% and ≥ 30% in both conditions, see **Supplemental information** for details).

We found that the mean pRF eccentricity of the included voxels was significantly smaller in the high-SF mapping condition (mean ± S.E.M.: high-SF: 4.41° ± 0.16°; low-SF: 4.88° ± 0.16°, paired *t*-test: *t*_17_ = -6.50, *p* < 10^-5^, Cohen’s *d* = 1.533). Interestingly, however, no significant difference in the mean pRF size was found between the two SF mapping conditions (mean ± S.E.M.: high-SF: 0.98° ± 0.04°; low-SF: 1.00° ± 0.04°, paired *t*-test: *t*_17_ = -0.69, *p* = 0.502) (**Figure 3d**). One possibility is that the difference in pRF size between the two conditions varied with eccentricity, which was concealed to the averaged results. To test this possibility, we fitted a line relating pRF eccentricity with pRF size for each SF mapping condition for each individual subject. We found that pRF size increased with pRF eccentricity in a highly similar rate between the two SF mapping conditions (mean ± S.E.M.: high-SF mapping: slope: 0.11 ± 0.01, intercept: 0.62 ± 0.05; low-SF: slope: 0.12 ± 0.01, intercept: 0.61 ± 0.06) with no significant difference in either slope or intercept of the fitted lines (Wilcoxon signed-rank test: slope: *p* = 0.528; intercept: *p* = 0.811) (**Figure 3e**). These findings showed that the pRFs were more clustered around the fovea under the high-SF stimulation, while SF channel enhancement was not likely to have a major influence on pRF size.

It should be noted that the findings reported above could result from the pRF displacement that occurred at the global scale or in certain parts of the visual space. To arbitrate between these two possibilities, we investigated the consistency of pRF displacements in different sub-regions of the visual space, and the pRF density difference between the two SF mapping conditions. Because the pRF profiles in all four quadrants of the visual field showed a similar pattern, the data were collapsed across both horizontal and vertical meridians into a single quadrant to maximize sensitivity (Ho & Schwarzkopf, 2022). In the first analysis, we partitioned the combined quadrant with a 6-by-6 grid and binned the voxel-wise pRFs in each SF mapping condition with respect to the grid, which yielded two binning schemes. For each binning scheme, a displacement vector was computed to characterize the local pRF displacement (from the low-to the high-SF mapping condition) in each grid cell. Because the binned data in either scheme contained noise and not a perfect reflection of the true pRF positions (Stoll et al., 2022), the two weighted displacement vectors were averaged to reduce condition-specific noise and to control for statistical bias (Altan et al., 2024). We found that the averaged displacement vector pointed to the fovea in almost every grid cell (**Figure 3f**). To statistically evaluate this observation, we calculated the angular deviation between the average displacement vector and the direction of the fovea in each grid cell (see **Methods** for details). As such, a deviation value of 0° indicates a displacement towards the fovea, while a deviation of 180° indicates a displacement away from the fovea. As shown in **Figure 3g**, the distribution of angular deviations across different grid cells centered around 0° (mean phase: -0.38°; *V*-test: *v* = 25.20, *p* < 10^-8^). In the second analysis, we reconstructed a continuous map of pRF density by calculating the moving average of the voxel-wise pRF locations for each SF mapping condition, and computed the difference between the two maps (**Figure 3h**). Interestingly, we observed a clear gradient from the periphery to the fovea in the difference map from which two oppositely signed clusters emerged (non-parametric cluster-based permutation test: *p* < 0.05, multiple comparisons corrected). While the positive cluster indicated concentration of pRFs in the vicinity of the fovea under the high-SF stimulation (cluster peak eccentricity: 0.59°), the negative cluster indicated dispersion of pRFs under the low-SF stimulation in the periphery (cluster peak eccentricity: 8.43°). Together, these observations converged on a global-scale displacement of V1 pRFs elicited by different SF mapping stimuli. Specifically, the high-SF mapping stimulus led to a more concentrated tiling of pRFs around the fovea.

### SF channel modulation connects pRF displacement with perceptual distortion

Although our imaging results demonstrated a global displacement of V1 pRFs elicited by SF channel modulation, it remained unknown whether and how the pRF displacements were functionally linked to the perceptual distortion. To bridge between the perceptual and the neural findings, we modeled the population SF response based on the behavioral data in the out-scanner experiment, and measured the SF channel modulation effect based on the model. The behaviorally measured SF channel modulation effect was then used to predict the pRF changes. The population SF response elicited by each checkerboard background was modeled as the weighted sum of individual SF channels that were characterized as a bank of logarithmic Gaussian functions peaked at different SFs (0.2 to 1.7 cycles/° in steps of 0.25 cycles/°). The channel weight parameters encode the underlying contribution of each SF channel, and thus a change of channel weight reflects the enhancement or the inhibition of the corresponding channel (**Figure 4a**).

**Figure 4.**
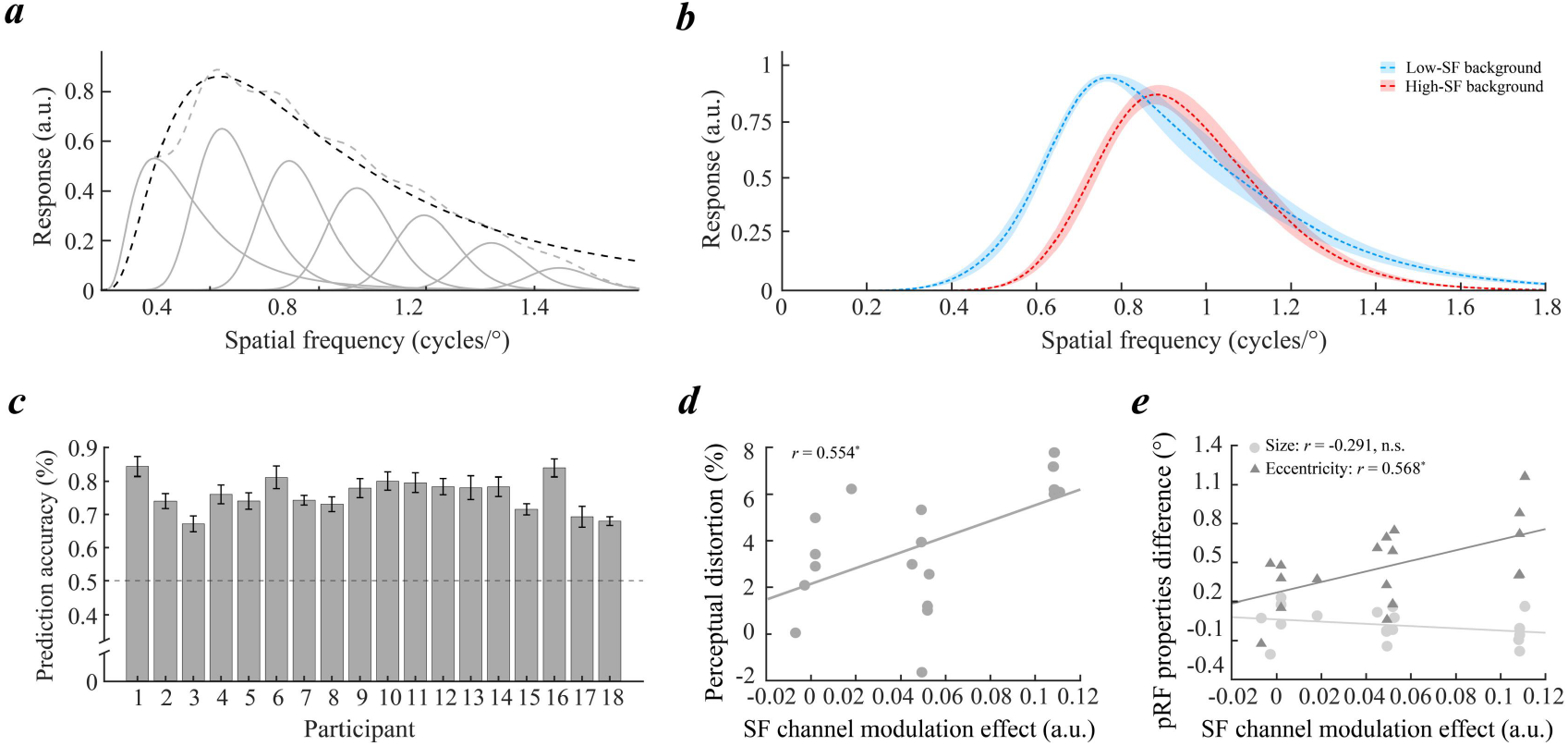
Computational modeling. ***a***. Illustration of the population SF response (the black dashed curve) modeled as the logarithmic Gaussian function fitted to the weighted sum of different SF channels (the gray dashed curve). Gray solid curves denote the idealized tuning functions of the SF channels. ***b***. Averaged population SF response corresponding to the high-(red) and the low-SF (blue) backgrounds across all subjects. The model of population SF response was fitted individually based on each subject’s behavioral data in the out-scanner experiment. Shaded areas denote one S.E.M. ***c***. Eight-fold cross-validated accuracy of the model-based prediction of the trial-by-trial behavioral response for each individual subject. Error bars denoted one S.E.M. across the cross-validation folds. The black dotted line indicates chance-level performance. ***d***. Correlation between the SF channel modulation (high-SF background – low-SF background) and the magnitude of perceptual distortion. ***e***. Correlation between the SF channel modulation and the difference in mean pRF eccentricity and pRF size between the two SF mapping conditions (low-SF mapping – high-SF mapping). *: *p* < 0.05, ***: *p* < 0.001, n.s.: not significant.

Subject’s trial-by-trial response was predicted by comparing the perceived length of the standard stimulus and the probe in each trial. The perceived length was calculated as the number of ruler graduations spanned by the bar, which was obtained by multiplying its physical length by the peak of the population SF response. Using a reverse correlation approach, we obtained the population SF response for each checkerboard background (Fernández et al., 2022).

We took two approaches to examine the validity of our model. First, we assessed its biological plausibility by reconstructing and comparing the population SF responses elicited by the two checkerboard backgrounds (**Figure 4b**). We found a significant difference in the peak of the population SF response (high-SF background: 0.83 ± 0.01 cycles/°, low-SF background: 0.78 ± 0.01 cycles/°, paired *t*-test: *t*_17_ = 4.98, *p* < 0.001, Cohen’s *d* = 1.174), as one would expect from a neural population sensitive to different SF stimulations. Second, we evaluated the prediction performance of the model by comparing the model prediction with the empirical responses across different trials for each individual subject. As shown in **Figure 4c**, the cross-validated prediction accuracy significantly exceeded the chance-level performance (50%) by a large margin for each and every subject (mean accuracy ± S.E.M. = 76.07 ± 1.22%, permutation test: *p* < 0.001, Bonferroni corrected for multiple comparisons), indicating that our model was able to predict subjects’ trial-by-trial responses with exceptional accuracy.

Having demonstrated the validity of our model, we then investigated whether the pRF changes could be predicted from the SF channel modulation effect measured by the model. For each subject, SF channel modulation effect was measured as the difference in the peak of the population SF response between the high- and the low-SF background conditions. We found a significant correlation between the perceptual distortion and the magnitude of SF channel modulation effect (Pearson’s *r* = 0.55, *p* = 0.017), which further supported the validity of the SF channel modulation measurement (**Figure 4d**). To quantify the pRF changes, we computed the difference in both mean pRF eccentricity and pRF size between the low- and the high-SF mapping conditions. As a result, the magnitude of SF channel modulation predicted the pRF eccentricity difference (*r* = 0.57, *p* = 0.014) but not the pRF size difference (*r* = -0.29, *p* = 0.242), indicating that modulation of SF channels was functionally coupled with pRF position displacements but had little or none influence on pRF sizes (**Figure 4e**). Taken together, these results suggested a three-way connection between SF channel modulation, distortion of perceived separation and pRF displacement in V1.

## Discussion

In the current study, we combined the use of psychophysics, fMRI and computational modeling to investigate the neural correlates of the putative mental ruler in spatial vision. We found that selective inhibition of different SF channels via psychophysical adaptation altered the perceived separation. In contrast, driving these SF channels led to an opposite effect, which was paralleled by a global-scale displacement of the population receptive fields (pRF) in V1. Using computational modeling, we further showed that both the perceptual distortion and the V1 pRF displacement could be predicted by the changes in population SF response. Our results unraveled an adjustable mental ruler represented by the bank of differently-tuned SF channels.

Our finding of an adjustable mental ruler based on SF channels aligns with the elegant theory of an internal visual metric that weaves the ‘fabrics’ of visual space. As the division of the mental ruler affords a common magnitude system (i.e. visual metric) for spatial property perception, our findings bear on several classical geometrical illusions. For example, in Ebbinghaus-like figures, the apparent size of the central object can be inflated, independently of other factors, by decreasing the size or the proximity of the surrounding contextual objects (Kirsch & Kunde, 2021; Roberts et al., 2005; Urale & Schwarzkopf, 2023). In Ponzo-like illusions, visual objects located among the aggregated local image elements (e.g. multiple converging lines that form the linear perspective) or embedded within dense texture patterns are typically perceived larger even when other available pictorial depth cues remain unchanged (Kirsch & Kunde, 2024; Yildiz et al., 2022). Common in these scenarios is the local dominance of high-SF signals in the image region containing the target object, which leads to a finer division of the mental ruler. As a result, the perceived size is inflated as the target object spans more ruler graduations. In addition, there is also an excellent agreement between the Oppel-Kundt illusion and our idea. In this illusion, a part of the visual field subdivided by an array of dense visual elements appears larger than the undivided part of the same spatial extent, as the intervening elements introduces a strong signal consistent with the mental ruler with fine division (Kundt, 1863; Landwehr, 2021; Wade et al., 2017). Moreover, our findings are reminiscent of the functional division of labor between global and local processing. According to our idea, abrupt local changes in luminance (i.e. edges) are optimally resolved by the mental ruler with a fine division signaled by high-SF channels, while the continuous, global-scale luminance variations are best captured by the mental ruler with a coarse division signaled by low-SF channels. Tipping the balance between the high- and the low-SF channels adjusts the mental ruler’s division, which facilitates the processing of either global or local stimulus features in different tasks (Flevaris et al., 2010, 2011, 2014; Flevaris & Robertson, 2016). Together, our findings offer a novel unified mechanistic account for a plethora of visual phenomena that were originally thought to be mediated by different mechanisms.

Moreover, our findings are suggestive of a unified perceptual scaling system that does not dependent on attention. While pioneering studies of attention have shown that attention induces changes in both individual neuronal RFs (Connor et al., 1996, 1997; Womelsdorf et al., 2006, 2008) and SF sensitivity (Fernández et al., 2022; Jigo et al., 2021; Ramirez et al., 2025), both of which parallel the attentional effects in spatial property perception (Fortenbaugh et al., 2011; Gobell & Carrasco, 2005; Suzuki & Cavanagh, 1997), so far there is no direct evidence in support of such a three-way connection outside the domain of attention. Our findings filled this gap by showing that SF channel modulation was quantitatively related to both global pRF displacements and distortions in perceived separation in the absence of attentional modulation. As a further note, concentration of neuronal RFs is thought to promote an attention-centered spatial code in which the locations of all visual stimuli are represented relative to the attentional focus (Connor, 2006). This might explain the RF shifts centered at the task-relevant object in previous studies (He et al., 2015; Ni et al., 2014), and the pRF displacement in relation to fixation observed in our study. Interestingly, however, we did not observe any systematic changes in pRF size between different SF mapping conditions, nor did we find any quantitative relationship between pRF size and perceived separation. Since the ability of integrating sensory information of a visual neuron is largely dependent on the size of its RF (Moutsiana et al., 2016; Song et al., 2013, 2015), a change in RF size might therefore reflect an adjustment in the spatial range over which sensory information is integrated. However, such spatial range adjustment was unlikely to be involved in the pRF mapping, which might be a reason for the absence of pRF size changes in our case. Another potential reason is that the SF range we investigated might be too narrow in terms of detecting a change in pRF size. Nevertheless, our findings demonstrate that SF channel weighting is a critical limiting factor of the neuronal RF topography regardless of whether attention is involved.

Although our neural findings primarily concern the pRF displacements in V1, this is not to say, however, that all forms of perceptual distortions are solely mediated by the earliest cortical processing stage. In natural viewing environments, the retinal images projected by visual objects are transformed to retain object constancy in accordance with the contextual cues to visual depth and 3-D surface configuration. These cues are extracted and processed in the cortical network comprised of higher-order extrastriate visual areas and parietal–occipital junctions (Altan et al., 2025; Nasr & Tootell, 2018; Ng et al., 2021; Tsao et al., 2003). Moreover, it was also found that the spacing between neighboring visual objects is influenced by the contextual motion, which is processed in hV5/MT+ (Özkan et al., 2021; Ross et al., 1997; Zirnsak et al., 2014). These findings hence suggest that the scaling of spatial property information in complex visual scenes might be accomplished in areas beyond early visual cortex. In this sense, we tentatively proposed a recurrent mechanism that centers at V1. On one hand, processing of the incoming visuospatial signals is constrained by the mental ruler signaled by the population SF response, which is recalibrated by top-down feedbacks. On the other hand, reweighting of the local SF channels might facilitate the interpretation of visual contexts in a way that is more consistent with the mental ruler, as reflected in the common technique of inducing depth perception on 2-D images by visual artists (Gregory, 1997). Within this ongoing recurrent process, V1 serves as the critical pivot where the final percept is directly read out from the topography of its neural activities that carry the richest spatial information at the highest spatial resolution among all the cortical visual areas.

In summary, we provide, for the first time, the neural evidence for the putative mental ruler in spatial vision. By connecting SF channel modulation, V1 pRF displacements and distortions in spatial property perception, our findings unravel a unified cognitive scaling mechanism that commonly governs the perception of spatial property information in different contexts.

## Methods

### Subjects

A total of 37 students (22 females, 18-26 years old) from South China Normal University participated in the study. There were 22 participants in the psychophysical adaptation experiment (two subjects completed both tasks), and there were 18 participants in both the out-scanner experiment and the ensuing fMRI scanning (three of them also participated in the psychophysical adaptation experiment). All participants had normal or corrected-to-normal vision and reported no known neurological or psychiatric disorders. All participants were naïve to the purpose of the study. Written informed consents were collected individually before the experiments. The experimental protocol was approved by the human subject review committee in South China Normal University (Ethics Approval: SCNUPSY-2021-342; IORG Registration: IORG0011738).

### Psychophysical adaptation experiment

Visual stimuli were generated using Psychophysics Toolbox 3 (http://psychtoolbox.org/) and were displayed on an LCD monitor (refresh rate: 60 Hz, resolution: 1920×1080 pixels, background luminance: 13.44 cd/m^2^). The adaptors were two full-contrast, phase-randomized sinusoidal gratings subtending an area of 10°×10°, which were simultaneously presented in the two visual fields. The centers of the adaptors aligned with the fixation on the horizontal meridian at an eccentricity of 10°. The spatial frequency of one adaptor (the fixed adaptor) was fixed at 0.25 cycles/°, while the SF of the other adaptor (the matching adaptor) could be one of the four SF values (0.25, 0.75, 1 and 1.5 cycles/°). Both adaptors had the same orientation, and were drifting in synchrony at a constant speed of 2.5°/s with their directions reversed every 5 seconds. For the dot-pair task, the task stimuli were two pairs of horizontally aligned dots (dot size: 0.4°). For the shape task, the task stimuli were two ellipses with vertical major axes (length: 8.75°). The eccentricities of the task stimuli were the same as that of the adaptors. In both tasks, the task stimulus ipsilateral to the fixed adaptor served as the standard stimulus (gap: 3.5°; aspect ratio: 5/2), while the task stimulus contralateral to the fixed adaptor served as the probe that varied in the gap of the dot-pair or the aspect ratio of the ellipse.

The psychophysical adaptation experiment was carried out in a behavioral test room. Subjects viewed the stimuli at a distance of 80cm with their heads rested on a chin rest. Each subject completed a total of 12 blocks for each of the four matching adaptors. In half of the blocks, the fixed adaptor was presented in the left visual field. Each block began with a 30-second pre-adaptation. In a trial, the task stimuli were presented for 100ms immediately after a 5-second topping-up adaptation. Subjects performed a two-alternative forced choice (AFC) task in which they indicated which of the two dot-pairs appeared more separated (the dot-pair task) or which of the two ellipses appeared more elongated (the shape task). A staircase procedure was employed in which the dot-pair separation or the aspect ratio of the probe was continuously adjusted on a trial-by-trial basis. For each observer, the behavioral data was fitted with a Logistic function weighted by the number of trials for each dot-pair separation or aspect ratio using PSignifit 3.0 (Fründ et al., 2011), which yielded the psychometric curves describing the percentage of the trials in which the probe was chosen over the standard stimulus. The point of subjective equality (PSE) of the standard stimulus was obtained as the point corresponding to 50% probability from the psychometric curve. For each matching adaptor, the magnitude of the adaptation effect was measured as the relative difference between the perceived and the physical property (gap or aspect ratio) of the standard stimuli in percentage units:

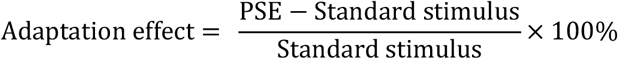

### Out-scanner experiment

The setup of the out-scanner experiment was the same as that of the psychophysical adaptation experiment. The task stimuli were two oblique yellow bars (bar width: 0.35°) presented on the background of two full contrast, phase-randomized checkerboards of different SFs (1.5 and 0.25 cycles/°). Both checkerboard backgrounds subtended an area of 6°×6°, which were located at an eccentricity of 5° on the horizontal meridian. The bar superimposed on the low-SF checkerboard served as the standard stimulus (bar length: 3.5°), while the bar on the high-SF background was the probe with variable length. Notably, the standard stimulus along with the low-SF checkerboard background could appear in either visual field with equal probability. The checkerboard backgrounds were presented for a total duration of 2000ms, during which the two checkerboards were solely presented for 1900ms before the bar stimuli was presented on top of them for 100ms. To minimize potential adaptation or temporal expectation effects, the polarity of the checkerboard stimuli was reversed with 50% probability every 166ms. Subjects were required to indicate which of the two bars appeared longer. Next, two Gaussian white noise patches were presented for 2s after a 1.025s blank screen to restore the default, unmodulated SF channel responses. The white noise patches were matched with the background stimuli in mean luminance and size. Similarly, a staircase procedure was employed to find the PSE of the standard stimulus by adjusting the length of the probe on the high-SF background. Each subject completed a total of 8 staircase blocks. We quantified the perceptual distortion effect as the relative difference between the physical length of the standard stimulus and the PSE in percentage units.

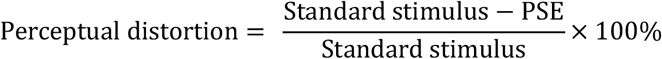

### pRF mapping

MRI data was acquired with a 3T Siemens MAGNETOM Prisma scanner in the MRI center of South China Normal University with a 20-channel phase-array head coil. Visual stimuli were back-projected via a video projector (refresh rate: 60 Hz, resolution: 1024×768) onto a screen positioned inside the scanning bore. Subjects viewed the stimuli through a mirror mounted on the head coil. The viewing distance was 80 cm. BOLD signals were measured using an echo planar imaging sequence (TE: 33 ms, TR: 2000 ms, FOV: 192×192 mm^2^, flip angle: 90°). 25 coronal slices orthogonal to the calcarine sulcus were collected in each functional volume, which encompassed the occipital lobe and a small portion of the cerebellum (slice thickness: 2 mm, gap: 0 mm). A T1-weighted high-resolution 3D structural image was acquired at the beginning of the scanning using the 3D-MPRAGE sequence.

The functional scanning was comprised of 4 separate runs for retinotopic mapping and 8 runs for pRF model estimation. In the retinotopic mapping runs, subjects were presented with a rotating wedge and an expanding/contracting ring that created traveling waves of neural activity in visual cortex. The stimuli for pRF mapping had two SFs (high-SF: 1.5 cycles/°; low-SF: 0.25 cycles/°). For each SF mapping condition, the hemodynamic response function (HRF) in V1 was first measured for each subject in a single run containing 12 trials. In each trial, a full-contrast flickering checkerboard stimulus subtending an area of 16°-by-16° centered at fixation point was presented for 2s, which was then followed by a 30-s blank interval. The HRF was obtained by fitting the convolution of a 6-parameter double-gamma function with a 2-s boxcar function to the BOLD response elicited by the checkerboard. PRFs were mapped with a full-contrast flickering, phase-randomized checkered bar (width: 2°) oriented either horizontally or vertically, which moved in two opposite directions (up-down, left-right). The bar swept through the visual field in 16 steps with a duration of 2s for each step. Each pRF mapping run consisted of 4 sweeps, with every two sweeps followed by a 12-s blank screen. There were three pRF mapping runs for each SF condition. Throughout the entire scanning session, subjects performed a color change detection task on the central fixation point to maintain fixation and to control for attention.

Functional MRI data was preprocessed using SPM 12 (https://www.fil.ion.ucl.ac.uk/spm/software/spm12/) and custom scripts written in MATLAB 2019b (The MathWorks). The first 6s of the BOLD signals were discarded to minimize transient magnetic saturation effects. The functional volumes were corrected for head motion and were co-registered with the anatomical scan. No subjects exhibited excessive head movement (>1 mm in translation or >1° in rotation). The voxel time series were band-pass filtered between 0.015 and 0.1 Hz to remove low-frequency noise and to improve pRF model fitting efficiency. Retinotopic area V1 was delineated on the reconstructed cortical surface using FreeSurfer (https://surfer.nmr.mgh.harvard.edu/) using the standard phase-encoded method (Engel et al., 1997). For both the high- and the low-SF pRF mapping stimuli, voxel-wise pRF model parameters were estimated using the same coarse-to-fine search method described in Dumoulin and Wandell (2008). For each voxel, the effective stimulation at each time sample was calculated as the spatial overlap between the mapping stimulus and the pRF model. The model has three parameters (x_0_, y_0_ and σ), which describe the location and the size of the pRF, respectively. The predicted BOLD signal time series was then obtained by convolving the HRF with the effective stimulation time course. Model parameters were adjusted to obtain the best fit to the empirical BOLD signal. Only the voxels for which the pRF model could explain at least 25% of the raw data variance in both SF mapping conditions were included for further analyses.

### pRF model analyses

We computed the mean pRF eccentricity and the mean pRF size of all the included voxels corresponding to the two SF mapping conditions for each subject. Each index was then statistically compared across all subjects between the two conditions. Moreover, in order to quantify the relationship between the pRF size and the pRF eccentricity, we first binned the voxel-wise pRFs in eccentricity steps of 0.2°, and calculated the average pRF size in each eccentricity bin for each subject. A line was fitted to each individual subject’s data. We then compared the slope and the intercept of the fitted lines between the two SF mapping conditions using the non-parametric Wilcoxon signed rank test. To obtain the confidence interval (C.I.) of the size-eccentricity relationship at the group level, a bootstrapping method was employed in which we iteratively sampled the subject’s data with replacement and calculated the average of the slope and intercept in each iteration (Mo et al., 2018). This procedure was repeated for 1000 times, yielding a bootstrapped distribution of slope and intercept, respectively. The 95% C.I. of the slope and the intercept were determined from the 97.5 and the 2.5 percentiles of the corresponding distribution.

We conducted two analyses to investigate the local pRF displacement pertaining to different parts of the visual field. In both analyses, the voxel-wise pRF data from all subjects were pooled together and were merged into one quadrant (i.e. the first quadrant) in prior. To collapse the data from different quadrants, we converted the pRF centers located in the second, the third and the fourth quadrants to their symmetrical points about the vertical meridian, the fovea and the horizontal meridian, respectively. In the first analysis, we partitioned the merged quadrant into a 6-by-6 grid and binned the voxel-wise pRFs in each SF mapping condition, which yielded two binning schemes.

For each binning scheme, we identified the voxels whose pRF centers were located within each grid cell, and computed a displacement vector from the mean pRF coordinate in the low-SF mapping condition to that in the high-SF mapping condition. The two displacement vectors were weighted by the percentage of the voxels corresponding to that cell, and were combined to obtain the average displacement vector for each cell. To characterize the direction of local pRF displacements in each cell, we calculated the angular deviation between the average displacement vector and the direction from the average (across the two binning schemes) of the mean pRF coordinate in the low-SF mapping condition to the fovea. Statistical evaluation of the angular deviation was conducted using the circular *V* test (Berens, 2009). In the second analysis, we first obtained a spatial map of pRF locations for each SF mapping condition, where each point of the map was indexed by the number of the voxel-wise pRFs centered at that point. We then reconstructed a continuous map of the pRF density by calculating the moving average of the pRF location map using a 3.5°-by-3.5° aperture. Finally, we computed the difference in the pRF density map between the two SF mapping conditions. Statistical assessment of the pRF density difference was conducted using the non-parametric cluster-based permutation analysis (Maris & Oostenveld, 2007). To define the supra-threshold clusters, we obtained the point-by-point cluster-defining threshold for each location using a randomization procedure (Wolff et al., 2017). In each iteration, we exchanged the pRF data between the two SF mapping conditions in a random subset of subjects and performed the same analysis procedure as described above. The randomization process was repeated for 1000 times to generate a null distribution of the pRF density difference at each location and we chose the 95th percentile of the null distribution as the cluster-defining threshold for both the empirical and the surrogate data. We identified the supra-threshold cluster with the maximal absolute value in the surrogate pRF density map in each iteration, which yielded the null distribution of cluster-based values from which the statistical threshold (*p* < 0.05, corrected for multiple comparisons) for clusters was obtained.

### Computational modeling

We modeled the population SF response as the weighted sum of multiple differently-tuned SF channels. The model was parameterized by three vectors (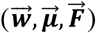) that described the weight, the preferred SF and the tuning width of each SF channel, respectively:

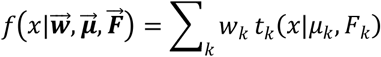

where *x* denotes to the stimulus SF. *W*_*k*_ represents the contribution of the *k*-th SF channel to the population response (i.e. channel weight). The idealized tuning function of the *k* -th SF channel was modeled as a logarithmic Gaussian function *t*_*k*_ (*x*), with the preferred SF and the full width at half maxima (FWHM) denoted as μ_*k*_ and *F*_*k*_, respectively. We considered seven SF channels centered at 0.2 to 1.7 cycles/° in steps of 0.25 cycles/°, hence the idealized tuning function is given as:

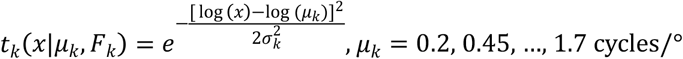

where σ_*k*_ is computed from μ_*k*_ and *F*_*k*_ as:

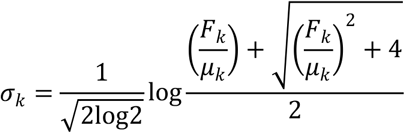

A reverse correlation approach was used to find the vector of channel weights and the vector of tuning widths (i.e. FWHM) corresponding to each checkerboard background condition. First, we obtained the population SF response by fitting a logarithmic Gaussian function to the weighted sum of SF channel tuning functions for each checkerboard background condition. Second, we computed the perceived length of each bar by multiplying its physical length by the preferred SF of the corresponding population response. Third, a prediction of subject’s response was derived in each trial by numerically comparing the perceived length of the two bars. Finally, we obtained the parameters that yielded the best fit to the behavioral data by minimizing the summed error between the predicted and the empirical trial-by-trial responses.

## Data availability statement

Data and scripts for generating the plot shown in this article are publicly available at https://osf.io/68pmy/?view_only=d87a888ca6144f1ca3d432294a80a152.

## Author contributions

S.Z., C.M., and F.F. designed the experiment. S.Z. performed the experiment. L.M. supervised the study. C.M., S.Z., and F.F. analyzed the data. S.Z., C.M. and F.F. wrote the paper.

## Acknowledgments

This study was supported by National Science and Technology Innovation 2030 Major Program (2022ZD0204802) to F.F., National Natural Science Foundation of China (32471104 to C.M. and T2421004, 31930053 to F.F.), Opening Project of Key Laboratory of Brain, Cognition and Education Sciences (South China Normal University), Ministry of Education, to L.M., and Basic and Applied Research Foundation of Guangzhou, China (SL2023A04J00644) to C.M.

