## Supplementary material for "Spatial frequency channels mediate a mental ruler for spatial vision in humans"

**Supplementary information**

To reduce potential concerns of idiosyncrasy in voxel selection, we performed the same analyses involving the pRF data as reported in the main text using a more lenient and a more stringent criterion (> 20% and >30% variance explained by the pRF model). The mean percentage of the included voxels was 56.16% (S.E.M.: 2.05%) for the former and 28.45% (S.E.M.: 1.98%) for the latter, respectively. Both analyses yielded highly consistent results with respect to the results reported in the main text, which are summarized in **Figure S1** & **S2**.

The average pRF eccentricity was significantly smaller in the high-SF mapping condition in comparison to the low-SF condition (20% explained variance: mean ± S.E.M.: high-SF: 4.72° ± 0.14°; low-SF: 5.22° ± 0.14°, paired *t*-test: *t*_17_ = -6.59, *p* < 10^-5^; 30% explained variance: mean ± S.E.M.: high-SF: 4.20° ± 0.18°; low-SF: 4.61° ± 0.18°, paired *t*-test: *t*_17_ = -5.06, *p* < 10^-4^), while there was no significant difference in pRF size between the two SF mapping conditions (20% explained variance: mean ± S.E.M.: high-SF: 0.95° ± 0.04°; low-SF: 0.97° ± 0.04°, paired *t*-test: *t*_17_ = -0.78, *p* = 0.445; 30% explained variance: mean ± S.E.M.: high-SF: 1.01° ± 0.04°; low-SF: 1.02° ± 0.05°, paired *t*-test: *t*_17_ = -0.45, *p* = 0.658) (**Figure S1a** & **S2a**). Moreover, pRF size increased with eccentricity at a similar rate between the two conditions (20% explained variance: mean ± S.E.M.: high-SF: slope: 0.104 ± 0.011, intercept: 0.609 ± 0.043; low-SF: slope: 0.101 ± 0.009, intercept: 0.633 ± 0.064, **Figure S1b**; 30% explained variance: mean ± S.E.M.: high-SF: slope: 0.122 ± 0.012, intercept: 0.615 ± 0.063; low-SF: slope: 0.134 ± 0.011, intercept: 0.578 ± 0.054, **Figure S2b**), with no significant difference between the slopes or intercepts (Wilcoxon signed-rank test: 20% explained variance: slope: *p* = 0.557; intercept: *p* = 0.913; 30% explained variance: slope: *p* = 0.306; intercept: *p* = 0.446).

For the analyses of pRF displacement between the two conditions, the averaged positional change vectors in different grid cells were found consistently pointing towards the fovea (20% explained variance: mean deviation angle: 3.56°; *V*-test: *v* = 26.14, *p* < 10^-9^, **Figure S1 c&d**; 30% explained variance: mean deviation angle: 0.35°; *V*-test: *v* = 19.48, *p* < 10^-5^, **Figure S2 c&d**), which were in line with our reports in the main text. Moreover, the pRF density difference between the two conditions also showed a similar pattern (20% explained variance: foveal cluster peak eccentricity: 0.24°, peripheral cluster peak eccentricity: 8.87°; 30% explained variance: foveal cluster peak eccentricity: 0.77°, peripheral cluster peak eccentricity: 8.87°) (**Figure S1e** & **S2e**). Finally, SF channel modulation predicted the pRF eccentricity difference (20% explained variance: *r* = 0.58, *p* = 0.013; 30% explained variance: *r* = 0.56, *p* = 0.016) but not the pRF size difference (20% explained variance: *r* = -0.27, *p* = 0.282; 30% explained variance: *r* = -0.20, *p* = 0.435) (**Figure S1f** & **S2f**).

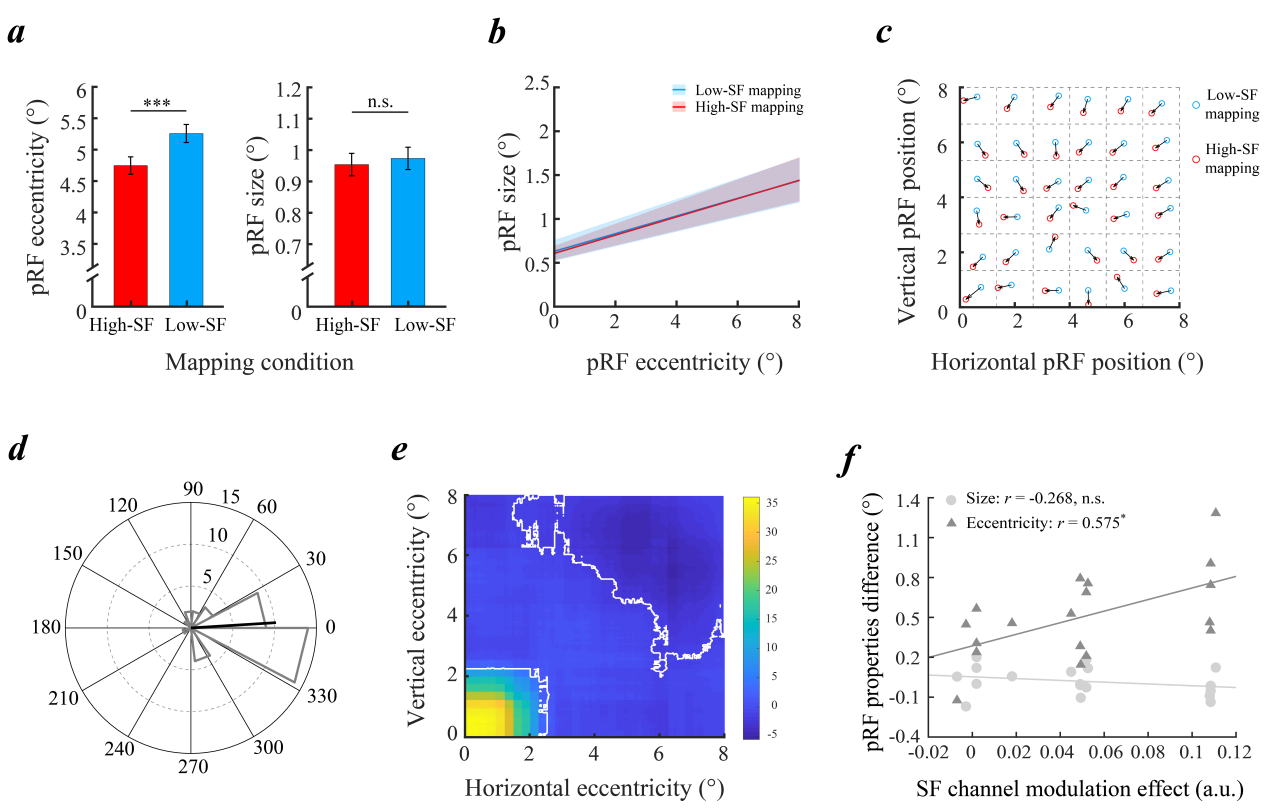

**Figure S1. Results of the pRF data analyses based on a more lenient criterion (>20% variance explained)**

***a*)** Comparison of the pRF eccentricity and the pRF size. ***b*)** Relationship between pRF size and pRF eccentricity. ***c*)** Positional change vectors pertaining to different parts of the visual space. ***d*)** Distribution of the angular deviation between the positional change vector and the direction from the empirical cell center to the fovea. ***e*)** Spatial map of the pRF density difference between the two SF mapping conditions. ***f*)** Correlation between the magnitude of SF channel modulation effect and the difference in pRF size and pRF eccentricity. Notations, symbols and color codes are identical to those reported in the main text (see **Figure 3d-3h & Figure 4e** for details)

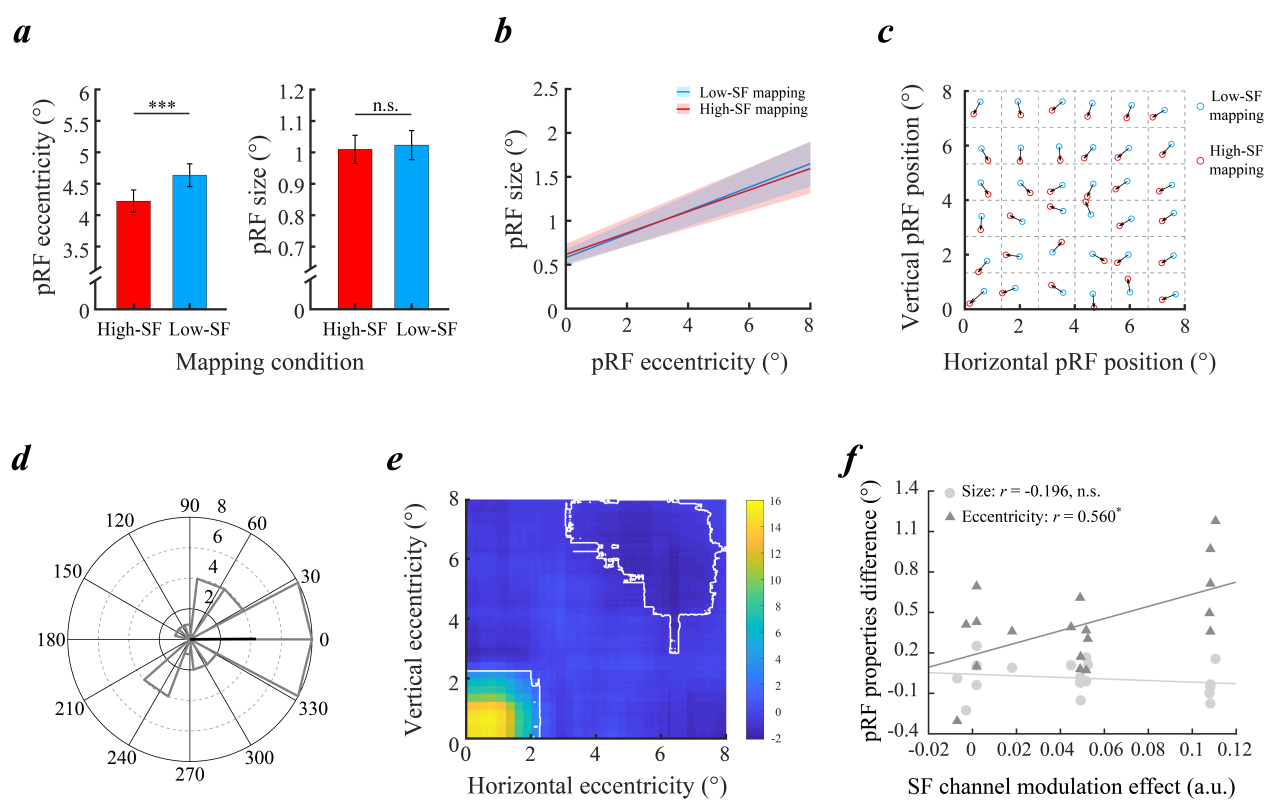

**Figure S2. Results of the pRF data analyses based on a more stringent criterion (>30% variance explained)**

Data is presented in the same way as in **Figure S1**. Notations, symbols and color codes are identical to those reported in the main text (see **Figure 3d-3h & Figure 4e** for details)
